# Tracking down the molecular architecture of the synaptonemal complex by expansion microscopy

**DOI:** 10.1101/821298

**Authors:** Fabian U. Zwettler, Marie-Christin Spindler, Sebastian Reinhard, Teresa Klein, Andreas Kurz, Markus Sauer, Ricardo Benavente

**Author notes:** These authors contributed equally to this work (Fabian U. Zwettler, Marie-Christin Spindler). Correspondence and requests for materials should be addressed to M.S. and R.B. ( and).

## Abstract

The synaptonemal complex (SC) is a meiosis-specific nuclear multiprotein complex that is essential for proper synapsis, recombination and segregation of homologous chromosomes. We combined structured illumination microscopy (SIM) with different ExM protocols including U-ExM, proExM, and magnified analysis of the proteome (MAP) to investigate the molecular organization of the SC. Comparison with structural data obtained by single-molecule localization microscopy of unexpanded SCs allowed us to investigate ultrastructure preservation of expanded SCs. For image analysis, we developed an automatic image processing software that enabled unbiased expansion factor determination. Here, MAP-SIM provided the best results and enabled reliable three-color super-resolution microscopy of the SCs of a whole set of chromosomes in a spermatocyte with 20-30 nm spatial resolution. Our data demonstrate that post-expansion labeling by MAP-SIM improves immunolabeling efficiency and allowed us thus to unravel previously hidden details of the molecular organization of SCs.

---

Imaging technologies are central platforms that drive fundamental research in virtually all disciplines across the biological and medical sciences. However, the diffraction barrier of classical fluorescence microscopy has hindered obtaining high-resolution information about the molecular architecture of protein assemblies and their interrelations. So far, only electron microscopy (EM) techniques provided a spatial resolution enabling the investigation of the molecular composition and structure of multiprotein complexes^1^. Super-resolution microscopy methods now can provide spatial resolution that is well below the diffraction limit of light microscopy approaching virtually molecular resolution^2, 3^. Physical expansion of the cellular structure of interest represents an alternative approach to bypass the diffraction limit and enables super-resolution imaging on standard fluorescence microscopes. For this purpose, expansion microscopy (ExM) has been developed and successfully applied to visualize cellular structures with ∼ 70 nm lateral resolution by confocal laser scanning microscopy^4^.

The original ExM protocol used functionalized antibody-oligonucleotide conjugates that bind to target proteins and cross-link covalently into a swellable hydrogel during polymerization. After degradation of native proteins by enzymatic proteolysis, the sample expands ∼4.5-fold in water^4^. To circumvent fluorophore loss during polymerization and protease digestion alternative ExM protocols have been introduced enabling imaging of proteins, RNA, and bacteria in cultured cells, neurons, and tissues also in combination with super-resolution microscopy^5–11^. For example, protein-retention ExM (ProExM)^6^ and magnified analysis of the proteome (MAP)^7^ have been developed that cross-link proteins themselves into the polymer matrix. Replacing protein digestion by heat and chemical induced denaturation allows post-expansion immunolabeling of chemically embedded proteins. To further increase the achievable resolution, the expansion factor has been increased up to 20-fold^12, 13^. However, such high expansion factors dramatically reduce the labeling density and consequently also the achievable structural resolution and require ultimately single-molecule sensitive imaging methods to visualize such extremely diluted fluorescence signals. Furthermore, some doubts remained concerning uniform three-dimensional (3D) expansion and preservation of ultrastructural details especially of multiprotein complexes. Very recently, it has been shown that various expansion protocols do not completely preserve the 3D molecular architecture of centrioles. Only by careful optimization of the expansion protocol ultrastructural details of centrioles could be truthfully preserved by U-ExM^14^.

In the present study, we tested the suitability of different ExM protocols for investigation of the molecular architecture of mammalian synaptonemal complexes (SCs) by structured illumination microscopy (SIM). With EM^15–19^ and single-molecule localization microscopy^20^ data available about the distribution of SC proteins, it is ideally suited as a benchmark structure to evaluate isotropic expansion and structure preservation of different ExM protocols.

ExM-SIM has already been used to investigate the three-dimensional (3D) organization of *Drosophila* SCs with a lateral resolution of ∼ 30 nm^21, 22^. To locate expanded sample as close as possible above the coverslip, they had to be dehydrated, cryosectioned into 10 µm sections, and then again expanded and mounted on a coverslip. Finally, SIM with an oil-immersion objective and minimal spherical aberration has been performed. Here, we optimized sample handling to avoid dehydration and cryosectioning steps and enable super-resolution imaging of expanded SCs by SIM. Therefore, we used nuclear spreadings of mouse spermatocytes, a widely used technique to study nuclear proteins, specifically in the field of meiosis. The spreading of the SCs directly onto the surface of the coverslip results in the localization of the proteins close to the coverslip post-expansion and allows the use of a water-immersion objective for imaging of hydrogels without spherical aberrations. Further, we adapted the hydrogel composition that enabled us to completely detach and transfer the entire sample from the glass surface to the hydrogel. These optimizations and the use of SC spreadings enabled isotropic expansion of the sample and multicolor post-expansion epitope labeling with no need for dehydration and cryosectioning, thus providing a quicker, less error prone approach to study the 3D organization of nuclear proteins.

Very recently, ExM has been combined with single-molecule localization microscopy by STORM to elucidate the molecular organization of the murine chromosome axis of SCs on nuclear spreadings^23^. Using ExSTORM on 2.7x expanded samples, the authors achieved a lateral resolution of 20-30 nm, similar to the spatial resolution demonstrated previously by *d*STORM imaging of unexpanded SCs^20^. With a ∼3-4x expansion factor in combination with a 2-fold increase in spatial resolution provided by SIM, currently Ex-SIM represents the method of choice for 3D multicolor super-resolution imaging of multiprotein complexes such as SCs. Therefore, we developed a robust workflow for Ex-SIM on nuclear spreadings together with an automated image processing software (‘Line Profiler’) to simplify the implementation of multicolor Ex-SIM and refined data analysis. The developed method allowed us to unravel new details of the molecular organization of SCs.

## Results and Discussion

### Analyzing SIM images of expanded SCs

Synaptonemal complexes (SCs) are meiosis-specific multiprotein complexes that are essential for synapsis, recombination and segregation of homologous chromosomes, resulting in the generation of genetically diverse haploid gametes, the prerequisite for sexual reproduction^24, 25^. The SC exhibits an evolutionarily conserved ladder-like organization composed of two lateral elements (to which the chromatin of homologous chromosomes is associated) and a central region. The central region is formed by a central element running between the lateral elements, and numerous transverse filaments connecting the lateral elements and the central element (Fig. 1). Early EM 3D reconstructions show the ribbon-like lateral elements (LEs) of the SC spanning across the nucleus while turning around the own axis^18^. In mammals, eight SC protein components have been identified so far: the proteins SYCP2 and SYCP3 of the lateral elements, SYCP1 of transverse filaments and the proteins SYCE1, SYCE2, SYCE3, TEX12 and SIX6OS1 of the central element^24, 25^. The assembly of the SC proteins into an elaborate molecular architecture is hereby tightly coordinated with essential meiotic processes and therefore conserved across species^20–25^. Consequently, localization maps of SC proteins are required to unravel the function of the molecular architecture of the SC in synapsis and recombination and thereby the overall success of meiosis.

**Figure 1.**
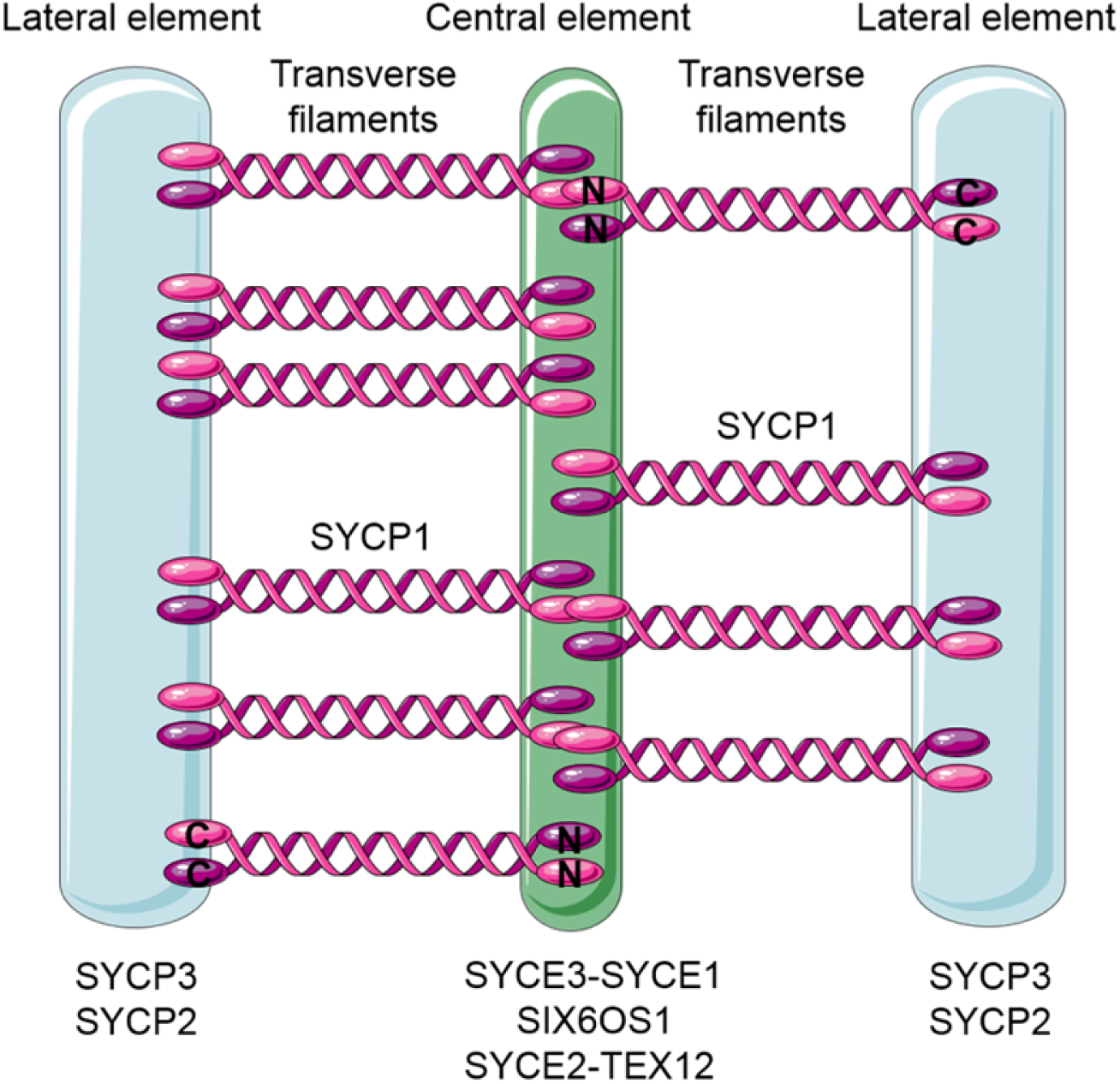
The synaptonemal complex. Schematic representation of the tripartite SC structure showing the two lateral elements (LEs) consisting of SYCP2 and SYCP3 flanking the central element comprised of SYCE1/2/3, Tex 12 and SIX6OS1. The transverse filament protein SYCP1 is connecting the lateral element and the central element with the SYCP1 C-terminus residing in the lateral and the N-terminus in the central element^34^.

In order to elucidate the precise molecular architecture of the SC, nanoscale resolution provided by either EM or super-resolution microscopy is required. Using immunolabeling and super-resolution microscopy by *d*STORM the position of different proteins of the SC on nuclear spreadings have been visualized with approximately 20-30 nm lateral resolution. The images revealed that the lateral element protein SYCP3 shows a bimodal distribution separated by 221.6 ± 6.1 nm (SD) (Fig. 2)^20^. This value is in accordance with distances measured between the centers of the two ribbons of parallel oriented lateral elements by EM^15–18^. With SC protein distribution data available, structure preservation and uniform expansion of different ExM protocols can now be efficiently evaluated.

**Figure 2.**
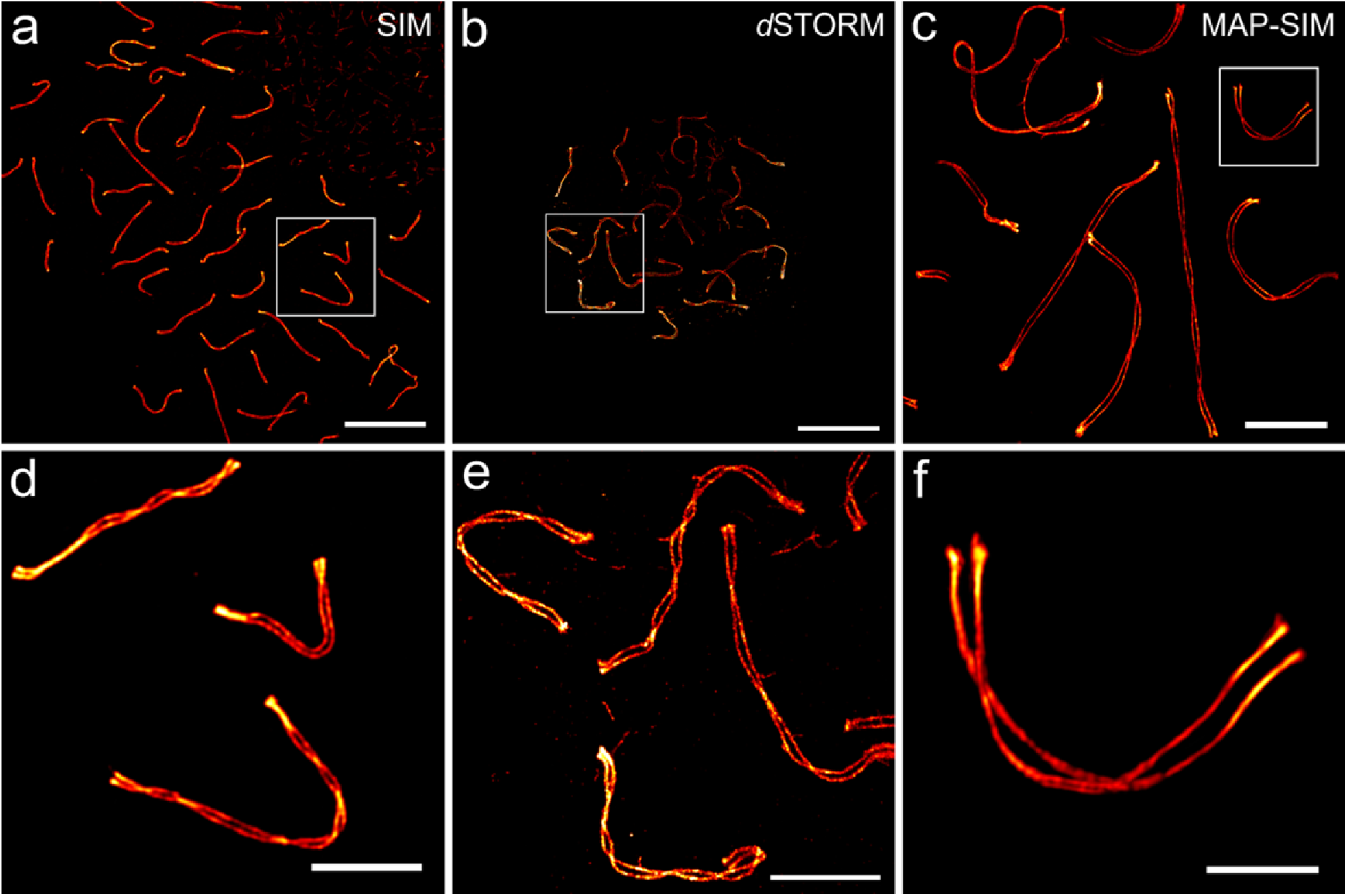
Super-resolution imaging of the lateral element protein SYCP3. **a,** SIM image of unexpanded SYCP3 labeled with Alexa Fluor 568. **b,** *d*STORM image of unexpanded SYCP3 labeled with Alexa Fluor 647. **c,** Expanded MAP-SIM SYCP3 signal (maximum intensity projection) labeled with SeTau-647. **d-f,** Magnified views of boxed regions in (**a**), (**b**) and (**c**), respectively. The images indicate that MAP-SIM provides a similar labeling density and spatial resolution as *d*STORM of 20-30 nm in the imaging plane. Scale bars. (**a**-**c**) 10 µm. (**d**-**f**) 3 µm.

We started with immunolabeling of SYCP3, the N-termini of SYCP1 (SYCP1N), and SYCE3 as proteins of the lateral element, the transverse filaments, and the central element of the SC, respectively on nuclear spreadings using three different ExM protocols. To automatically and objectively analyze the average position of fluorescently labeled SC proteins and determine distances between bimodal distributed proteins from cross-sectional profiles, we developed ‘Line Profiler’, an automated image processing software (https://line-profiler.readthedocs.io/en/latest/)^26^. Line Profiler uses several image-processing algorithms to evaluate potential regions of interest. In a first step all structures in the SYCE3 channel are reduced to lines with one pixel width by using a threshold and a skeletonize algorithm. The resulting pixel-coordinates are sorted and fitted with a c-spline. This gives an analytical description of the orientation of SYCE3 and therefore a good approximation for the center (line coordinates) and overall orientation of the helically arranged SYCP3 protein (2-channel mode) (Supplementary Fig. 1). Note that it is also possible to determine the orientation with a gradient image of the SYCP3 channel, if the SYCE3 channel cannot be evaluated (1-channel mode) (Supplementary Fig. 2).

To compute the distance between the SC strands, we applied a floodfill algorithm to the SYCP3 channel, leaving only areas embedded in closed shapes unequal to zero. In combination with a distance transform, i.e. a computation of the distance of each point within the area to the nearest point outside, the regions, where the helical structure of the SC is in plane (regions of maximal distance between the strands) are revealed. All line coordinates outside of these areas were discarded. A line profile is subsequently constructed for each remaining line coordinate perpendicular to the derivative of the c-spline. For averaging, the line profiles are post aligned to the center between their peaks. We determined the expansion factors of the different protocols by comparing the distances between the maximum intensities of the two SYCP3 strands of unexpanded and expanded SCs imaged by *d*STORM and SIM, respectively (Fig. 2). SYCP3 bimodal protein distributions of unexpanded SCs imaged with *d*STORM resulted in an average strand distance of 222 nm ± 33 nm (SD) consistent with our previous *d*STORM data^20^ (Supplementary Fig. 3).

### Optimization of SC expansion

We tested the pre-expansion labeling protocol proExM^6^, and the two post-expansion labeling protocols MAP^7^ and U-ExM^14^. Among the three ExM protocols tested MAP outperformed the other protocols resulting in an average peak-to-peak distance of cross-sectional profiles of the SYCP3 signals of 632 ± 73 nm (SD) determined using the 1-channel mode to analyze the cross-sectional profiles (**Methods** and Supplementary Fig. 2). With a bimodal distribution separated by 222 nm ± 33 nm (SD) as measured by *d*STORM from unexpanded SCs (Supplementary Fig. 3) the MAP protocol enabled three-color SIM imaging and provided an expansion factor of ∼ 2.9x. U-ExM enabled post-expansion labeling with various fluorophores and three-color SIM but expanded SCs showed structural breaks indicating insufficient incorporation of proteins into the gel matrix (Supplementary Fig. 4). U-ExM provided an expansion factor of ∼ 2.4x using the 1-channel mode method to analyze the cross-sectional profiles (Supplementary Fig. 5). On the other hand, proExM provided the largest expansion factor of ∼ 4.0x using the 1-channel mode but pre-expansion labeling resulted in a lower labeling density due to irreversible fluorophore destruction during free-radical polymerization (Supplementary Figs. 4 and 5)^4, 6^. Although the extent of irreversible fluorophore destruction during gelation varies across fluorophores multicolor imaging with pre-expansion labeling protocols remains challenging.

Furthermore, our MAP-SIM data (Fig. 3) show that the molecular architecture of the SCs is fully preserved demonstrating isotropic expansion. At first glance, this result appears surprising in light of our recent study evaluating the structure preservation of centrioles using different expansion protocols^14^. For centrioles, the MAP protocol was unsuited, as MAP-treated centrioles appeared even smaller when compared to unexpanded samples. Therefore, U-ExM has been introduced as a variation of the MAP protocol using weaker fixation that enabled excellent isotropic expansion and preservation of the centriole ultrastructure^14^. The main difference between centrioles and SCs is their biomolecular composition. While centrioles are solely composed of proteins, the SC consists of a tight association between DNA and proteins, which may affect expansion efficiency and isotropy. During fixation both proteins amongst each other as well as proteins and DNA are crosslinked. Possibly expansion of SCs requires stronger fixation conditions than used in the U-ExM protocol and at the same time an increased acrylamide concentration to preserve molecular identity and to enable detachment and full transfer of proteins into the gel. After gelation, the proteins are denatured at high temperatures using sodium dodecyl sulfate (SDS). Apart from denaturation, SDS also confers its negative charge to the proteins^27, 28^. As the DNA is negatively charged as well, crosslinked DNA and proteins repel each other. We hypothesize that the repulsion between DNA and proteins facilitates isotropic expansion of SCs when using the MAP protocol with stronger fixation conditions.

**Figure 3.**
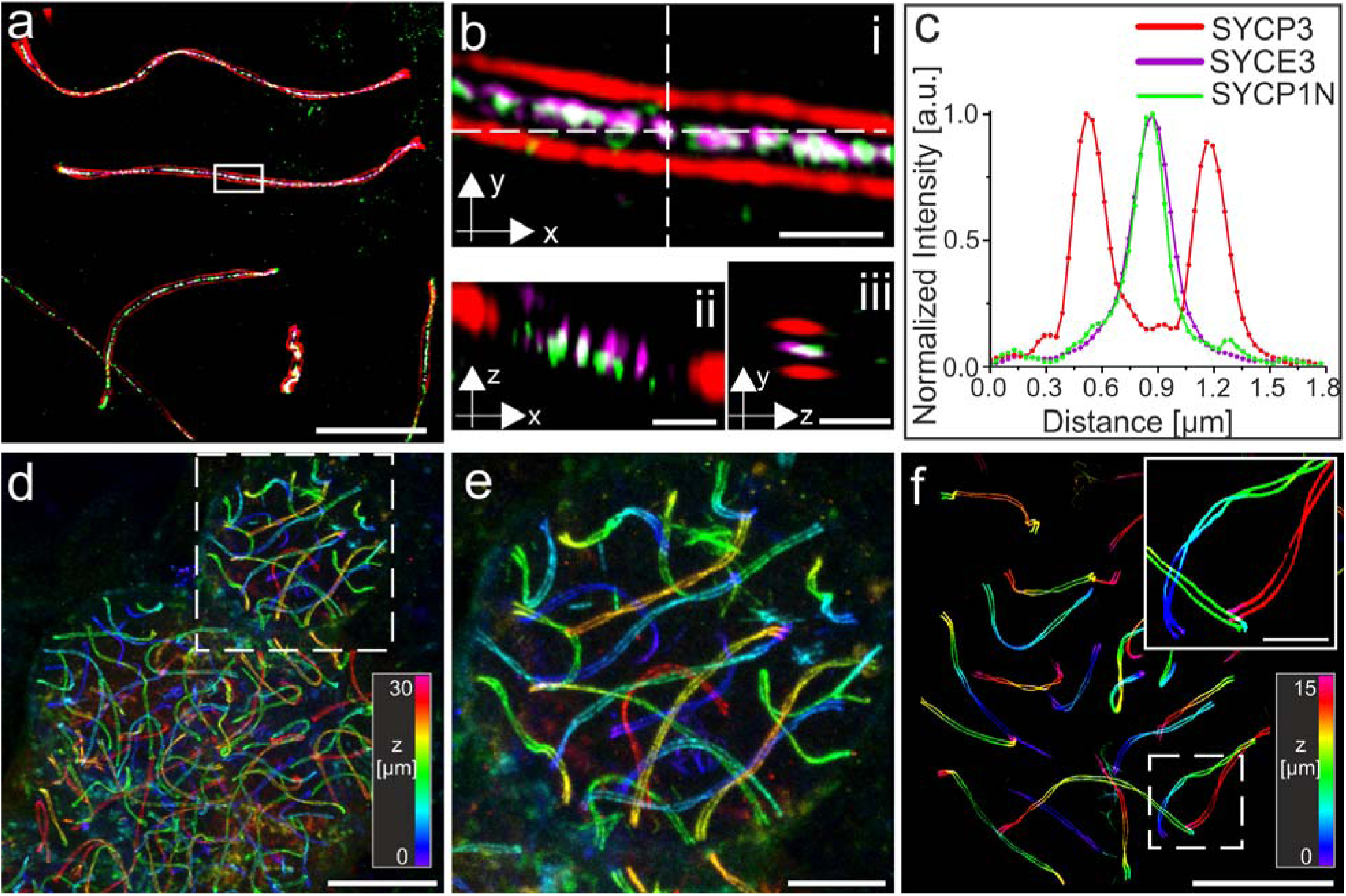
3D-Multicolor MAP-SIM of SYCP3, SYCP1 N-terminus, and SYCE3. **a,** SIM image of post-expansion SeTau-647 labeled SYCP3 as a component of the lateral element (red), the transverse filament SYCP1 N-terminus labeled with Alexa Fluor 488 (green) and SYCE3 of the central element labeled with Alexa Fluor 568 (magenta). **b,** Magnified views of white boxed region in (**a**). **ii,** Orthogonal view of horizontal white dashed line in (i). **iii,** Orthogonal view of vertical white dashed line in (i). **c,** Transversal intensity profile perpendicular to the orientation of the SC shown in (**b**). The selected section exhibits a bimodal distribution of the SYCP3 signal separated by 667.0 nm ± 7.1 nm. The SYCP1 N-terminus and SYCE3 signals of the section (b) show monomodal distributions with FWHM of 214.7 ± 6.9 nm and 258.3 ± 4.8 nm, respectively. **d**, Large field of view (100×100×30 µm^3^) 3D-Re-scan confocal microscopy image of spreaded cells. SYCP3 was labeled post-expansion with SeTau-647. **e,** Magnified view of boxed region in (d). **g,** 3D-MAP-SIM image of the SCs of an entire set of chromosomes in a spermatocyte visualized by post-expansion labeling of SYCP3 with SeTau-647. The inlet shows the enlarged view of the boxed region in (**f**). Scale bars. (**a**) 10 µm. (**b**-**d**) 1 µm. (**f**) 10 µm. (**g**) 15 µm. (**h**) 5 µm.

Overall, MAP-SIM with an optimized gel composition enables isotropic ∼ 2.9x expansion, efficient transfer of proteins into the hydrogel and molecular structure preservation of SCs. In combination with a two-fold resolution enhancement of SIM, MAP-SIM provides a similar spatial resolution as *d*STORM^20^ but in addition a higher immunolabeling efficiency because of the improved epitope accessibility of post-expansion protocols^14^. Therefore, we used our optimized MAP-SIM approach (Supplementary Fig. 6) in all following experiments to investigate details of the molecular architecture of SCs.

Previous *d*STORM experiments have shown that the C-terminus of SYCP1 localizes to the inner edge of the lateral element and the N-termini of SYCP1 interact in the central element^20^. These findings are in accordance with recent STORM experiments performed on 2.7x expanded nuclear spreadings^23^. Here, the SYCP1 N-terminus was located in the central element roughly 110 nm away from the SYCP3 labeled lateral element (LE) while the SYCP1 C-terminus localized 25 nm more inward to SYCP3, which corresponds to the inner edge of the lateral element^23^. Further, in *d*STORM experiments the width of the monomodal localization distributions of the N-terminus of transverse filament protein SYCP1 and the central element protein SYCE3 were determined to 39.8 ± 1.1 nm (SD) and 67.8 ± 2.1 nm (SD), respectively, in frontal views of the SC^20^. The broader signal distribution of SYCE3 localizations indicates that the interaction of SYCP1 and SYCE3 might not be limited to the N-terminus of SYCP1. This is consistent with expanded MAP-SIM protein distributions of 229.3 ± 1.2 nm (SD) (FWHM) for SYCE3 and 161.5 ± 1.3 nm (SD) (FWHM) for SYCP1N analyzed from frontal and lateral views of the SC (Supplementary Fig. 1).

Since *d*STORM requires efficient photoswitching of organic dyes in oxygen depleted thiol-buffer, it is currently limited to two-color experiments, whereby carbocyanine dyes such as Cy5 and Alexa Fluor 647 are the best suited fluorophores^29, 30^. In contrast, MAP-SIM provides similar resolution but enables imaging with up to three colors simultaneously without optimization of the photoswitching buffer conditions. The number of available laser lines thus only limits multicolor super-resolution microscopy experiments. Therefore, we immunolabeled the N-terminus of SYCP1, SYCE3, and SYCP3 with the same antibodies as used in the *d*STORM experiments in triple-localization MAP-SIM experiment (Fig. 3). The MAP-SIM images clearly exhibited similar details of the molecular architecture of SCs as single-molecule localization microscopy of unexpanded samples. In addition, the images confirmed isotropic expansion and preservation of the molecular structure of the SC. Intriguingly, spreading of SCs in combination with MAP allowed us to acquire 3D super-resolution images of large expanded samples, e.g. 100 x 100 x 30 µm^3^ (200 nm z-steps) by re-scan confocal microscopy (RCM)^31^ and 80 x 80 x 15 µm^3^ (110 nm z-steps) and larger by SIM, i.e. imaging of the SCs of a whole set of chromosomes in a spermatocyte with detailed structural information of single chromosomes (Fig. 3).

### MAP-SIM reveals a complex network organization of the SC central region

MAP-SIM images of lateral views of the SC reveal the topology of the transverse filament protein SYCP1 that is oriented perpendicularly to the lateral element protein SYCP3 and the central element protein SYCE3. *d*STORM and immunogold EM images showed a bimodal distribution of the N-terminus of SYCP1 and SYCE3 in twisted areas of the SC^19, 20^. EM tomography uncovered a multilayered organization of the central element in insects^32, 33^. On the other hand, EM tomography based 3D models of the murine SC recently revealed the absence of a layered organization of the SC central region^34^.

Therefore, we next analyzed the signal distribution of the N-terminus of the transverse filament protein SYCP1 by MAP-SIM of nuclear spreadings in more detail. In addition to analyzing the distribution of proteins in frontal views, we also generated line profiles of SC proteins from lateral views, i.e. twisted areas of the SYCP3 signal (Fig. 4 and **Supplementary Video 1**). In agreement with EM tomographic data^34^, the signal distributions of the MAP-SIM imaged SYCP1 N-terminus and SYCE3 indicated a far more complex distribution of the central region proteins than the bimodal distribution previously described by *d*STORM and immunogold EM^19, 20^.

**Figure 4.**
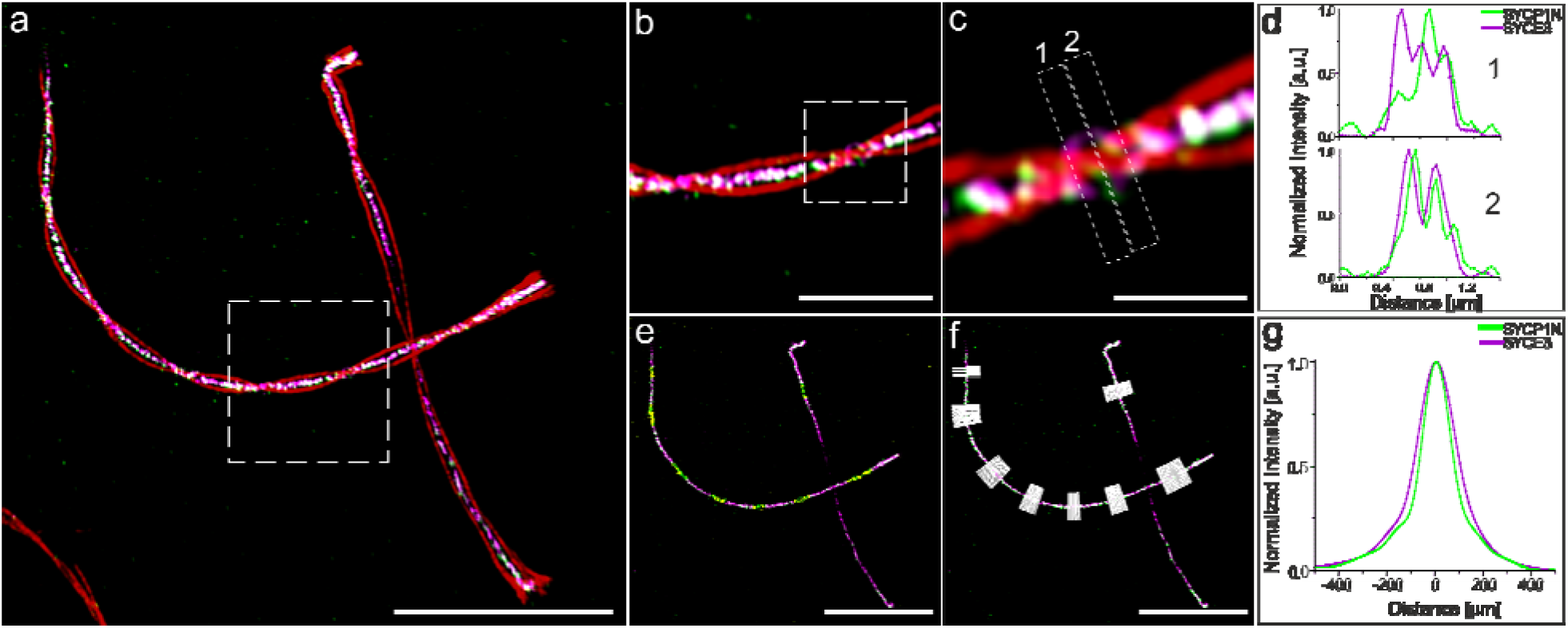
Protein distribution of the SYCP1 N-terminus reveals complex network organization of the SC central region. **a,** Lateral view MAP-SIM image of SYCE3 labeled with Alexa Fluor 568 (magenta), SYCP1N labeled with Alexa Fluor 488 (green) and SYCP3 labeled with SeTau-647 (red). Regions where the SC twists as visualized by the twisting SYCP3 signal (lateral view) show partially a multimodal organization of the N-terminus of the transverse filament protein SYCP1 and the central element protein SYCE3. **b,** Magnified view of white boxed region in (**a**). **c,** Enlarged view of highlighted region in (**b**) with two sites selected for protein distribution analysis (1, 2). **d,** Respective cross-sectional intensity profiles along SYCE3 and SYCP1N of the regions specified in (**c**). **e,** SYCE3 and SYCP1N signals as in (**a**) without SYCP3 signal. Frontal views of the two proteins used for cross-sectional profiles are shown in yellow. **f,** as in (**e**) with line profiles (white) perpendicular to the frontal view of the SYCE3 signal. **g,** Averaged intensity line profile of all profiles shown in (**f**) with a monomodal signal distribution of 168.0 ± 1.1 nm (SD) for SYCP1N and 211.43 ± 0.89 nm for SYCE3 derived from single Gaussian fitting. Scale bars, (**a**) 10 µm, (**b**) 3µm, (**c**) 1 µm, (**e**-**f**) 10µm.

In traditional immunofluorescence and immuno-EM preparations, antibodies have to compete for epitope accessibility of the densely packed central element. Free epitopes at the core of the central elements are difficult to access for IgG antibodies with a size of 10-15 nm taken that primary antibodies bound to the outer epitopes will shield more central epitopes. Using MAP-SIM epitope accessibility is improved due to the initial physical enlargement of the multiprotein complex before post-expansion labeling. The resulting higher labeling density, specifically at the core of the central element, provides a more fine-grained resolution of the molecular organization of the central element. Hence, MAP-SIM discloses new information about the molecular organization of the SC, particularly that the central element proteins are not organized unambiguously as distinct layers but they form a complex network composed of the transverse filament protein SYCP1 and the central element protein SYCE3 as well as other central element proteins in agreement with recent electron tomographic findings^34^.

### Molecular details of SC axes uncovered by MAP-SIM

One striking morphological feature of the lateral elements, which could not be visualized before by light microscopy, is their occasional splitting into two or more sub lateral elements (subLEs). Variations of subLEs have been observed across species from animals to plants to yeast in various EM preparations (histological sections, whole-mount preparations, spreadings)^15, 35–41^. Again, the continuous signal distribution of MAP-SIM allowed us to close the existing gap between light microscopy and EM and resolve the splitting of the two SYCP3 strands in murine pachytene spermatocytes (Fig. 5 and **Supplementary Video 2**). Spermatocytes contain 40 chromosomes that synapse as bivalents of homologous chromosomes along the synaptonemal complex in order to recombine. The fully synapsed bivalents hereby span the nuclear space with the two pairs of telomeres residing at distant sides of the nuclear envelope. In the XY pair, synapsis is confined to the pseudo-autosomal region and is therefore incomplete. In autosomes, MAP-SIM resolved a doubling of each of the SYCP3 strands (i.e. LEs) specifically, but not exclusively, at the sites where the helical synaptonemal complex twists (lateral view of the SC) and at the ends of the SYCP3 strands (Fig. 5 and **Supplementary Video 2**).

**Figure 5.**
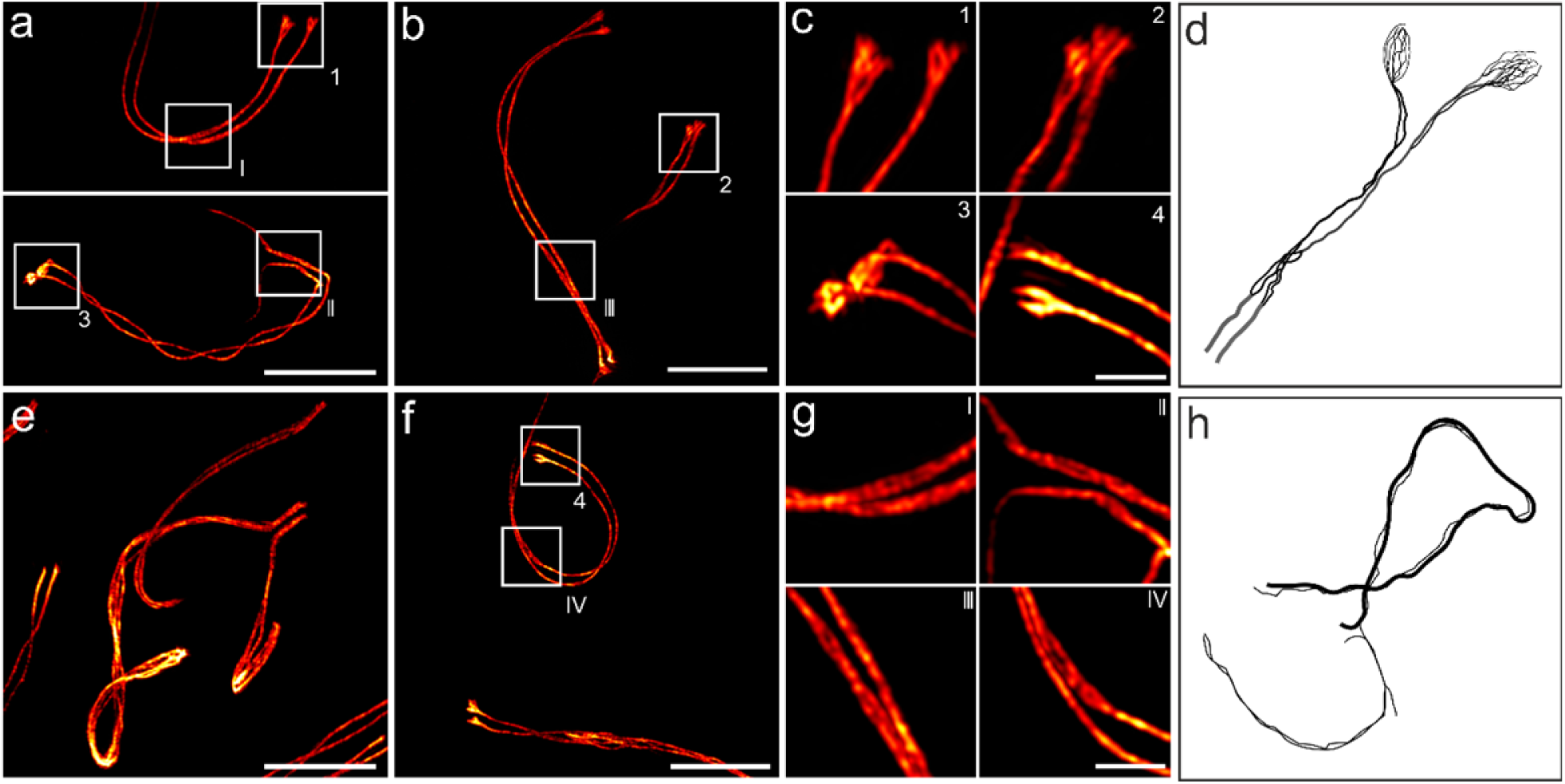
Structural details of the SC lateral element revealed by MAP-SIM of SYCP3. **a-c, f, g,** The SYCP3 signal shows occasional bifurcation along the two SC strands (I-IV) and various degrees of fraying at their ends depending on the respective pachytene stage (1-4). **e,** Unpaired regions of the XY pair also show a strong degree of fraying. **d, h,** Schematic representations of early EM reports of two or more sub lateral elements (subLEs) in mammals^38, 40^ in accordance with the splitting of the SYCP3 signal resolved by MAP-SIM in this study. **d**, Representation of the lateral element strand splitting in two and fraying at its end according to Figure 4b and 3a of del Mazo and Gil-Alberdi, 1986^38^. **h**, Fraying of LEs associated with unpaired regions of the XY pair modelled after Figure 10c of Dresser and Moses, 1980^40^. Scale bars, (**a-b**) 7 µm, (**c**) 1.5µm, (**e-f)** 7 µm, (**g**) 1.5µm.

With progression of prophase I, the degree of fraying at the end of the SYCP3 strands increases. In our experiments, we observed a bifurcation of the SYCP3 ends in mid-pachynema and a fraying of the ends into multiple strands in late pachynema/early diplonema (Fig. 5 and **Supplementary Video 2**). These observations are in agreement with EM findings of subLEs in murine spreadings^38^. Here, a multistranded organization of the lateral elements that arrange into two compact subLEs through interaction with the two sister chromatids has been proposed^37^. In EM images of silver-stained spreadings of mouse SCs, both axes appeared double or multistranded with a higher frequency of subLEs at unpaired regions^41^. Using MAP-SIM, we resolved for the first time the reported fraying of the axes into SYCP3 positive fibrils by light microscopy. In accordance with the conservation of the subLEs of the autosomes, also the fraying of the XY axes is common in mammals^42, 43^.

In mid-pachynema, the homologs are fully synapsed along the SC. Here, doubling of lateral elements is frequently observed and potentially related to the association with the two sister chromatids of the homologs. In diplonema, desynapsis starts and SCs gradually disassemble. At this stage, the lateral elements appear to disperse into individual SYCP3 fibrils. A strong degree of fraying has also been observed in unpaired regions of the XY pair. In 2014, Syrjänen *et al.* resolved the crystal structure of human SYCP3^44^. They showed that the tetrameric protein is approximately 20 nm long and is organized in an antiparallel arrangement that exposes its N-terminal DNA binding sites at either end. Binding to stretches of DNA, SYCP3 self-assembles into higher-order fibers that resemble the lateral elements in vitro. Based on doubling of the lateral elements observed in EM, it has been speculated that SYCP3 might assemble into one subLE per sister chromatid to prevent recombination between sister chromatids while maintaining chromatin cohesion^45^.

## Conclusion

To close the discussion about the preservation of molecular structures and uniformity of expansion, we performed a comparison of the 3D molecular architecture of mammalian SCs as visualized by unexpanded *d*STORM^20^ and expanded MAP-SIM. Taking advantage of the similar spatial resolution provided by the two methods, even smallest structural deviations are immediately apparent. We have shown that through isotropic structure preserving expansion combined with SIM, ultrastructural details of SCs can be revealed with standard immunofluorescence techniques in common sample preparations. Super-resolution microscopy techniques such as *d*STORM and PALM have enabled new insights into how proteins are organized in cells and multiprotein complexes, with a spatial resolution approaching virtually EM level^2, 3^. Nevertheless, structural details of the molecular architecture of multiprotein complexes remained largely accessible only to EM methods. This inability of super-resolution microscopy methods is often due to insufficient structural resolution, which is ultimately controlled by the labeling density. Our results demonstrate that post-expansion labeling protocols substantially increases the epitope accessibility for IgG antibodies resulting in higher labeling densities especially of sterically demanding multiprotein complexes^14^. Consequently, MAP-SIM provides improved ultrastructural resolution as demonstrated here for an important DNA-associated multiprotein complex.

## Online Methods

Animal care and experiments were conducted in accordance with the guidelines provided by the German Animal Welfare Act (German Ministry of Agriculture, Health and Economic Cooperation). Animal housing, breeding and were approved by the regulatory agency of the city of Wuerzburg (Reference 821-8760.00-10/90 approved 05.06.1992; according to §11/1 o.1 of the German Animal Welfare Act) and carried out following strict guidelines to ensure careful, consistent and ethical handling of mice.

### Reagents

Acrylamide (AA, 40%, A4058, Sigma), Acryloyl-X, SE, 6-((acryloyl)amino)hexanoic Acid, Succinimidyl Ester (A20770, Thermo Fisher), Agarose (A9539, Sigma), Ammonium persulfate (APS, A3678, Sigma), Bovine Serum Albumin (BSA, A2153, Sigma), Dimethyl sulfoxide (DMSO, D12345, Thermo Fisher), Ethanol (absolute, ≥ 99.8%, 32205, Sigma), Ethylenediaminetetraacetic acid (EDTA, E1644, Sigma), Formaldehyde (FA, 36.5-38%, F8775, Sigma), Guanidine hydrochloride (guanidine HCl, 50933, Sigma), Glucose (G8270, Sigma), Glucoseoxidase (Sigma), ß-Mercaptoethylamine (MEA, M9768, Sigma), N,N’-methylen-bisacrylamide (BIS, 2%, M1533, Sigma), N,N,N′,N′-Tetramethylethylenediamine (TEMED, T7024, Sigma), Normal goat serum (50197Z, Thermo Fisher), PBS (P5493, Sigma), Poly-D-lysine hydropromide (P6407, Sigma), Polyoxyethylene (20) sorbitan monolaurate solution (Tween-20, 10%, 93774, Sigma), Potassium hydroxide (P1767, Sigma), Proteinase K (P4850, Sigma), Sodium acrylate (SA, 97-99%, 408220, Sigma), Sodium chloride (NaCl, S7653, Sigma), Sodium dodecyl sulfate (SDS, L3771, Sigma), Tris base (T6066, Sigma), Triton X-100 Surfact-Amps Detergent Solution (10% (w/v), 28313, Thermo Fisher), Tween 20 Surfact-Amps Detergent Solution (Tween-20, 28320, Sigma).

### Murine spermatocyte cell spread preparation

Wildtype C57.6J/Bl6 mice were sacrificed using CO_2_, followed by cervical dislocation. Testes were resected and seminiferous tubules extracted and immersed in PBS after decapsulation of the testes. Next, nuclear spreadings were carried out as described by de Boer et al.^46^. Briefly, seminiferous tubules were transferred to hypotonic buffer and incubated for 60 minutes. Individual seminiferous tubules were transferred to sterile sucrose solution on a slide, disrupted with forceps and cells flushed out by resuspension with a 10 µl pipette. In parallel, poly-lysine slides were immersed in 1% formaldehyde, substituted with acrylamide in case of the MAP and U-ExM protocol (30% AA, 4% FA in PBS for MAP and 1% AA, 0.7 % FA in PBS for U-ExM). 20 µl of the testes cells in sucrose were transferred to a drop of the formaldehyde solution collected in a corner of the slide and spread evenly across the entire slide. Slides were incubated in a wet chamber for 2 hours and dried overnight in a wet chamber left ajar. To ensure ease of handling, nuclei were spread on round 18 mm coverslips (NO. 1.5H).

### Antibodies

Guinea pig and rabbit anti-SYCP1 (N-terminal amino acids 1-124)^2^, guinea pig anti-SYCP3 (N-terminal amino acids 27-38) and rabbit anti-SYCE3 (full length protein) were generated by Seqlab through immunizing the host with the respective peptides and affinity purified before use. Rabbit anti-SYCP3 (NB300-232; derived against the C-terminus) was purchased from Novus Biologicals and mouse anti-SYCP3 (ab97672; full length protein) was purchased from Abcam. Al647 goat anti-guinea pig IgG (H+L), highly cross-absorbed (A-21450) Invitrogen (ThermoFisher); goat anti-rabbit IgG (H+L) highly cross-adsorbed Alexa Fluor 568 (A-11036) was purchased from ThermoFisher; Alexa 488 conjugated AffiPure F(ab’)2 goat anti-guinea pig IgG (H+L) was purchased from Dianova (106-546-003). SeTau647 NHS (K9-4149; SETA Biomedicals) was conjugated to F(ab’)2 of goat anti-mouse IgG (SA-10225; ThermoFisher); F(ab’)2 goat anti-rabbit IgG (H+L) cross-adsorbed Alexa Fluor 647 (A-21246) and F(ab’)2 goat anti-mouse IgG (H+L) cross-adsorbed Alexa Fluor 488 (A-11017) were purchased from ThermoFisher. Supplementary Table 1 summarizes the immunolabeling used for the different experiments.

### Immunofluorescence of non-expanded SCs

Coverslips with fixated nuclear spreadings were washed with PBS. Unspecific epitopes were blocked for 1 hour in 10 % NGS. Nuclear spreadings were incubated face down on 100 µl of the primary antibody for 1 hour at room temperature in a humidified chamber. Spreadings were then washed with PBS and blocked for 30 minutes in 10 % NGS before incubation with the secondary antibody for 30 minutes at room temperature in a humidity chamber. For multi-color experiments, immunostaining was performed sequentially, starting with the antibody raised in mouse.

### Protein retention protocol (proExM)

#### Gel linker treatment

In order to cross-link amide groups into the polymeric hydrogel network, samples were incubated in freshly prepared amine reactive AcX solution (0.1 mg/ml) in PBS. Dessicated AcX stocks (10 mg/ml) stored as aliquots at −20°C were therefore resuspended in 10 µl anhydrous DMSO and diluted 1:100 in PBS (1x). Samples were then covered with 1 ml AcX solution per coverslip and incubated at room temperature in a humidified chamber overnight.

#### Gel formation, Digestion and Expansion

Hydrogel formation was carried out on a cell culture plate lid covered with parafilm. The plate was placed on ice to provide a cooled and flat hydrophobic gelation surface. 80 µl of pre-chilled (4°C) gelling solution (8.55 % SA, 2.5% AA, 0.15 % Bis-AA, 0.2 % APS, 0.2% TEMED, 11.7% NaCl, 1x PBS) were prepared from proExM Monomer stock solution consisting of a mixture of acrylic copolymers and crosslinking agent (8.55 % SA, 2.5% AA, 0.15 % Bis-AA, 11.7% NaCl, 1x PBS) that was supplemented with radical polymerization initiator APS and accelertator TEMED right before use. The gel solution was placed on parafilm and coverslips were put on top of the formed droplet with spread cells facing down. After 5min incubation on ice, crosslinking polymerization was allowed to occur for 1.5 hours at 37°C in a humidified chamber. Then samples were treated with 8 U /ml Proteinase K in Digestion Buffer (50 mM Tris pH (8.0), 1 mM EDTA, 0.5 % Triton X-100, 0.8 M guanidine HCl). For expansion of the samples, hydrogels were washed several times in double-deionized water until the maximum extent of swelling of the gels was reached.

### Magnified analysis of the proteome (MAP) and ultrastructure expansion microscopy (U-ExM) protocol

#### Gel linker treatment

In the case of MAP and U-ExM expanded samples, spreads were incubated in AA/FA solution (30% AA, 4% FA in PBS for MAP and 1% AA, 0.7 % FA in PBS for U-ExM) for 4 hours at 37°C in a humidified chamber before proceeding with gelation of the samples.

#### Gel formation, Denaturation and Expansion

Following AA/FA incubation, samples were washed three times for 5 minutes each in PBS (1x). Then polymerization of hydrogels was performed as described under ‘ProExM protocol’ using an optimized MAP Gel solution (7 % SA, 20 % AA, 0.05 % Bis-AA, 0.5% APS, 0.5% TEMED, 1x PBS) or U-ExM Gel solution (19% SA, 10% AA, 0.1% BIS-AA, 0.5% APS, 0.5% TEMED, 1x PBS). The MAP monomer solution composition was altered compared to the original recipe^7^ regarding the monomer to crosslinking agent ratio. After polymerization hydrogels were carefully removed from the coverslips and transferred directly into pre-heated (95°C) Denaturation buffer (200 mM SDS, 200mM NaCl, 50 mM Tris, pH 9.0) in 1.5 ml centrifuge tubes. Samples were then denaturated for 1 hour in a heating block incubator with closed tube lids. For swelling of the sample, hydrogels were washed with excess volume of double-deionized water that was exchanged several times until the maximum expansion level was reached.

### Post-expansion immunolabeling of MAP and U-ExM treated samples

For MAP and U-ExM treated samples immunostaining was performed post expansion. Fully expanded gels were incubated in Blocking buffer (0.15% BSA in PBS) twice for 30 minutes each. Gels shrink during this blocking step. Then primary antibodies were incubated sequentially for 3 hours each at 37°C in a humidified chamber with two 20 minutes washing steps in Blocking buffer following each antibody incubation step. Next a secondary antibodies mix was added simultaneously for 3 hours at 37°C in a humidified chamber. Samples were then washed twice for 30 minutes each with Washing Buffer (0.1 % Tween-20 in PBS) and once more overnight. Samples were then washed in double-deionized water and expanded back to the maximum hydrogel volume.

### Mounting of expanded samples

#### Poly-Lysine coating of cover glasses

24 mm round cover glasses (NO. 1.5. H) were sonicated in double-deionized water, 1 M KOH and absolute ethanol (≥ 99.8 %) for 15 minutes each. Following each sonication step glasses were rinsed with double-deionized water. After sonication glasses were finally dried at 100°C in an oven. Cover glasses were then covered with 0.1% Poly D-Lysine and incubated for 1 hour at room temperature. Next glasses were washed again with water and air-dried and stored in a closed glass container to avoid dust contamination. Coated coverslips were stored at 4°C and used for up to one week.

#### Fluorescent Marker treatment of cover glasses

Fluorescent beads introduced directly into the hydrogel show strong fluorescence loss caused by persulfate radicals during polymerization. For this reason we directly coated coverslips with fluorescent markers to perform channel alignment. Therefore, fluorescent marker stock suspension (0.1 µm, ∼1.8 × 10^11^ particles/mL, TetraSpeck Microspheres, Thermo Fisher) was vortexed for ∼1 minute and then diluted 1:1000 in PBS (1x) and vortexed again briefly. Glasses were covered with the suspension for 30 minutes at room temperature to let the fluorescent markers settle down. Then coverslips were washed with double-deionized water, air-dried and thereupon used for hydrogel immobilization with Agarose.

#### Agarose embedding

Expanded samples were cut into ∼1.5 x 1.5 cm pieces using a razor blade and excess water was removed carefully from the gels with laboratory wipes. Gels were then transferred onto Poly-Lysine coated coverslips. To further avoid drifting of the sample during long-term image acquisition gels were additionally embedded in 1 % (w/v) Agarose in water. Therefor a second uncoated round 18 mm-coverslip was placed on top of the cut hydrogel and melted Agarose (∼40°C) was carefully applied around the sides of the hydrogel using a pipette. Care was taken to avoid Agarose from flowing below the hydrogel. The agarose gel was subsequently hardened at 4°C for ∼10 minutes and the upper coverslip was removed carefully. Double deionized water was then added on the hydrogel to prevent dehydration during imaging.

### Imaging

#### Structured Illumination Microscopy (SIM) imaging

SIM imaging was performed using a Zeiss Elyra S.1 SIM imaging system consisting of an inverse Axio Observer. Z1 microscope equipped with a C-APOCHROMAT 63x (NA 1.2) water-immersion objective and four different excitation lasers (405 nm diode, 488 nm OPSL, 561 nm OPSL and 642 nm diode laser). Three-dimensional SIM imaging of the expanded sample was recorded on a PCO edge sCMOS (scientific complementary metal-oxide semiconductor) camera with 0.110 µm z steps adjusted by an inserted Z-Piezo stage. Three rotations of the grid pattern were projected on the image plane for each acquired channel. The red fluorophore was imaged before fluorophores with lower wavelengths to minimize photobleaching. Raw data images were processed using the Zeiss ZEN software (black edition).

#### Channel alignment for SIM images

Fluorescent beads mounted on the coverslip that were localized directly below the sample imaging area were set as lowest imaging plane and recorded with the sample for each 3D z-stack. For alignment and SIM processing images containing fluorescent markers were cropped out and used to align the channels in each recorded z-stack via the Zeiss Zen software channel alignment tool.

#### Rescanning confocal microscopy (RCM) imaging

RCM imaging was conducted on a Nikon TiE inverted microscope combined with an RCM unit (Confocal.nl). RCM derives from the image-scanning principle whereby pixel reassignment is carried out purely optomechanically^31, 47^. The unit is connected to a Cobolt Skyra (Cobolt, Hübner Group) laser unit providing four excitation laser lines (405 nm, 488 nm, 561 nm, and 640 nm). Images were recorded on an sCMOS camera (Zyla 4.2 P, Andor) using a 60x water-immersion objective (CFI Plan APO, 1.27-NA; Nikon) and a fixed 50 µm pinhole size. The setup was operated by the microscope software NIS-Elements (version 4.6).

#### dSTORM imaging

*d*STORM acquisition of unexpanded samples were conducted on a home-built widefield imaging setup as described previously^20^. The setup consists of an inverted IX71 microscope (Olympus) equipped with an oil-immersion objective (APON 60XOTIRF, NA 1.49, Olympus). For excitation of Alexa Fluor 647, a 639 nm OPS laser diode (Genesis MX639-1000 STM; Coherent) was used in quasi-TIRF mode. To avoid drift during acquisition a nose-piece stage (IX2-NPS, Olympus) is implemented in the microscope. Fluorescence light was collected onto an electron-multiplying charge-couple device (EM-CCD) camera (iXon Ultra 897, Andor). As photoswitching buffer 100 mM ß-Mercaptoethylamine in PBS (pH 7.4) supplemented with 10 % (w/v) glucose and 0.5 mg/ml glucose oxidase as oxygen scavenger system was used. For image reconstruction the ImageJ plugin ThunderSTORM^48^ was used.

### Protein position analysis

#### SC cross-sectional profiles using ‘Line Profiler’

The SYCE3 input image is convolved with a gaussian blur, compensating noise and intensity fluctuations. Via Otsu^47^ thresholding the image is converted into a binary image. Using Lee’s algorithm^7^ a skeletonize image is constructed reducing all expanded shapes to lines with 1 pixel width. Subsequently all connected pixels unequal to zero were sorted und checked for continuity, i.e. sharp edges mark a breakpoint initiating a new line. The pixel coordinates of each line were fitted with a c-spline. The c-splines coordinates and local derivatives are a good estimation for the center and orientations of the helix structure SYCP3 channel. Note that the orientation and center of the SYCP3 channel can also be estimated with Sobel filters, if the SYCE3 channel cannot be evaluated (‘1 channel’ mode). To receive the areas of maximum distance, i.e. the areas where the SYCP3 helix structure is in plane, a floodfill algorithm and a distance transform^8^, with a subsequent thresholding to 40 % of the maximum value, were applied. Using a logical and operation on the line coordinates and the thresholded distance transform image resulted in the desired source coordinates and directions for further evaluation. Line profiles were then constructed originating at the source coordinates perpendicular to the c-splines derivative or respectively in Sobel gradient direction at the edge points of the SYCP3 channel. The line profiles were post aligned at the center of the two global maxima. Average profiles were returned for each line and for the whole image.

#### Protein distribution of SYCE3 and SYCP1N

To determine the distribution of SYCE3, areas of the SYCE3 signal were manually chosen from regions where SYCP3 strands showed a helix crossing. These regions can be regarded as frontal views of the SYCE3 and SYCP1N proteins and were used for cross-sectional profiling (Fig 4).

#### Expansion factor determination

The expansion factor was determined by dividing the distances of the two SYCP3 strands measured from expanded SIM and unexpanded *d*STORM measurements. Here, strand distances of unexpanded SCs visualized by *d*STORM were determined using the ‘Line Profiler’ with Sobel filter settings for ‘1 channel’ analysis as described above and shown in Supplementary Fig. 3. Although the analysis of expanded MAP-SIM and U-ExM SIM data are possible using the SYCE3 channel to determine SC orientation, for reasons of comparability the expanded strand distance values were determined in the same ‘1-channel mode’ Line Profiler settings as for unexpanded *d*STORM analysis. Note that the ‘2-channel mode’ also includes areas which are closer to the SYCP3 strand crossings in comparison to the ‘1-channel mode’ (compare Supplementary Fig. 1 and Supplementary Fig. 2). The expansion factor was then calculated by dividing the resulting mean values of expanded SYCP3 distances with the distance mean value of unexpanded *d*STORM data.

#### 3D visualization of MAP-SIM data

3D MAP SIM data shown in **Supplementary Video 1** and **Supplementary Video 2** were visualized using the microscopy image analysis software IMARIS (version 8.4.1, Bitplane).

#### Averaging of cross-sectional profiles

The resulting cross-sectional profiles were finally averaged using the data analysis software Origin(Pro) Version 2016.

## Acknowledgements

The authors thank Dominic Helmerich for technical assistance with the SeTau647 labeled antibody. This work was supported by the German Research Foundation (TRR 166 Receptor Light project A04, and Be1168/8-1). M.C.S was supported by a Boehringer Ingelheim Fonds Travel Grant and a HHMI Grant to attend the 2018 CSHL course ‘Quantitative Imaging: From Acquisition to Analysis’ and wishes to thank the faculty of the course for excellent training and illuminating discussion benefitting the preparation of this manuscript-specifically Jennifer Waters, Talley Lambert, Hunter Elliot and Suliana Manley, as well as Marcelo Cicconet, Jessica Hornick, Anna Jost, and Michael Weber.

## Author contributions

F.U.Z., M.C.S., M.S., and R.B. conceived and designed the project. M.S and R.B. supervised the project. F.U.Z, M.C.S., A.K. and T.K performed all experiments. S.R. developed Line profiler. S.R. and F.U.Z. performed data analysis. All authors wrote and revised the final manuscript.

## Additional information

Supplementary information accompanies this paper at …….

## Ethics statement

Animal care and experiments were conducted in accordance with the guidelines provided by the German Animal Welfare Act (German Ministry of Agriculture, Health and Economic Cooperation). Animal housing, breeding and experimental protocols were approved by the regulatory agency of the city of Wuerzburg (Reference 821-8760.00-10/90; according to §11/1 No.1 of the German Animal Welfare Act) and carried out following strict guidelines to ensure careful, consistent and ethical handling of mice.

## Supplementary Figures

**Supplementary Table 1:**
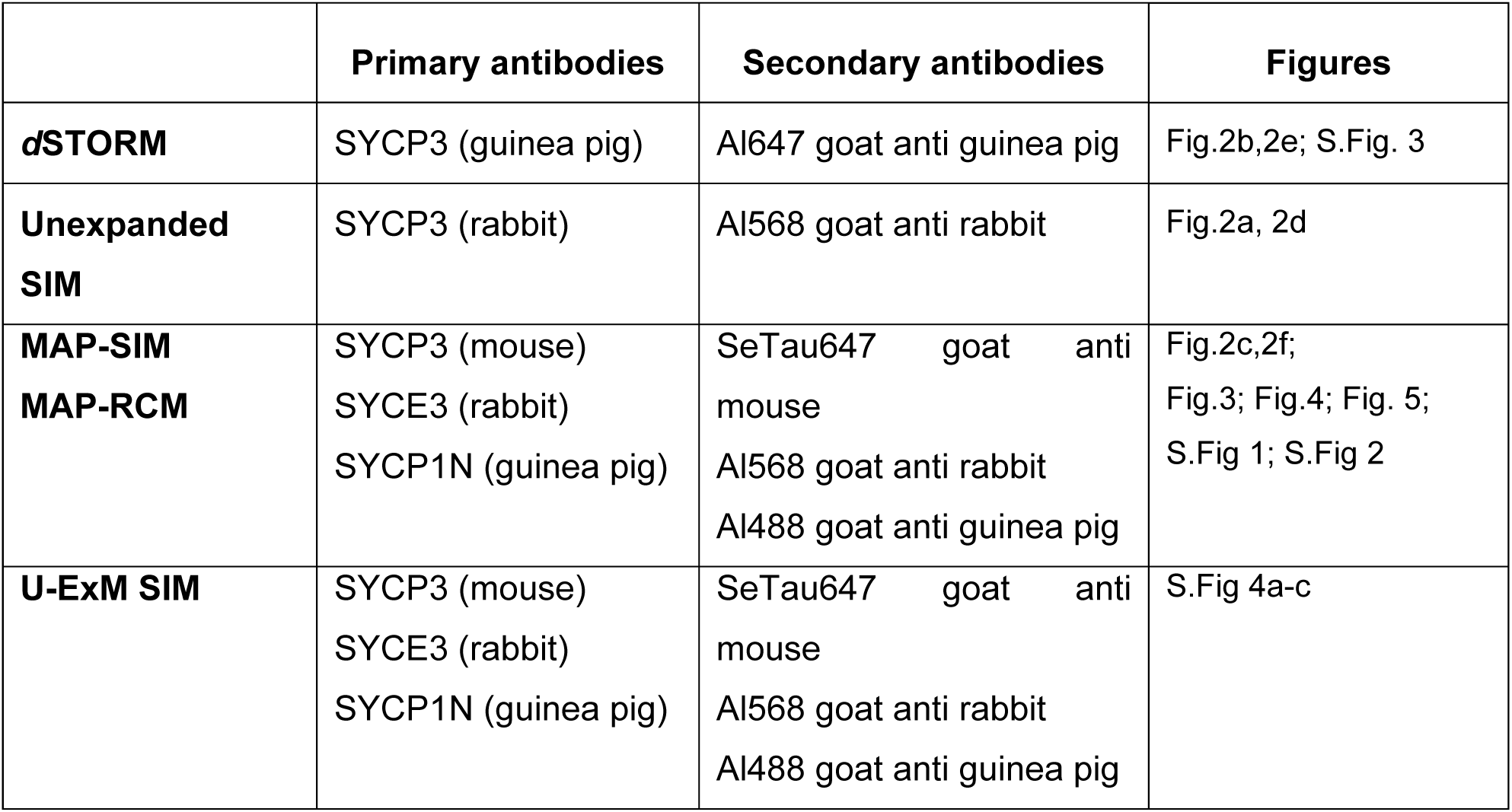
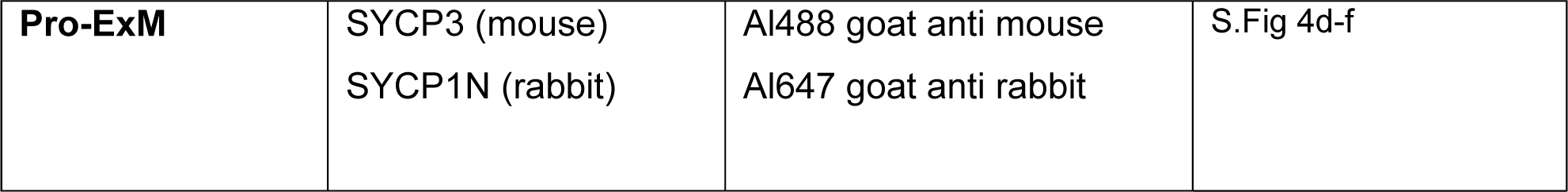
Immunolabeling used in the different experiments

**Supplementary Figure 1.**
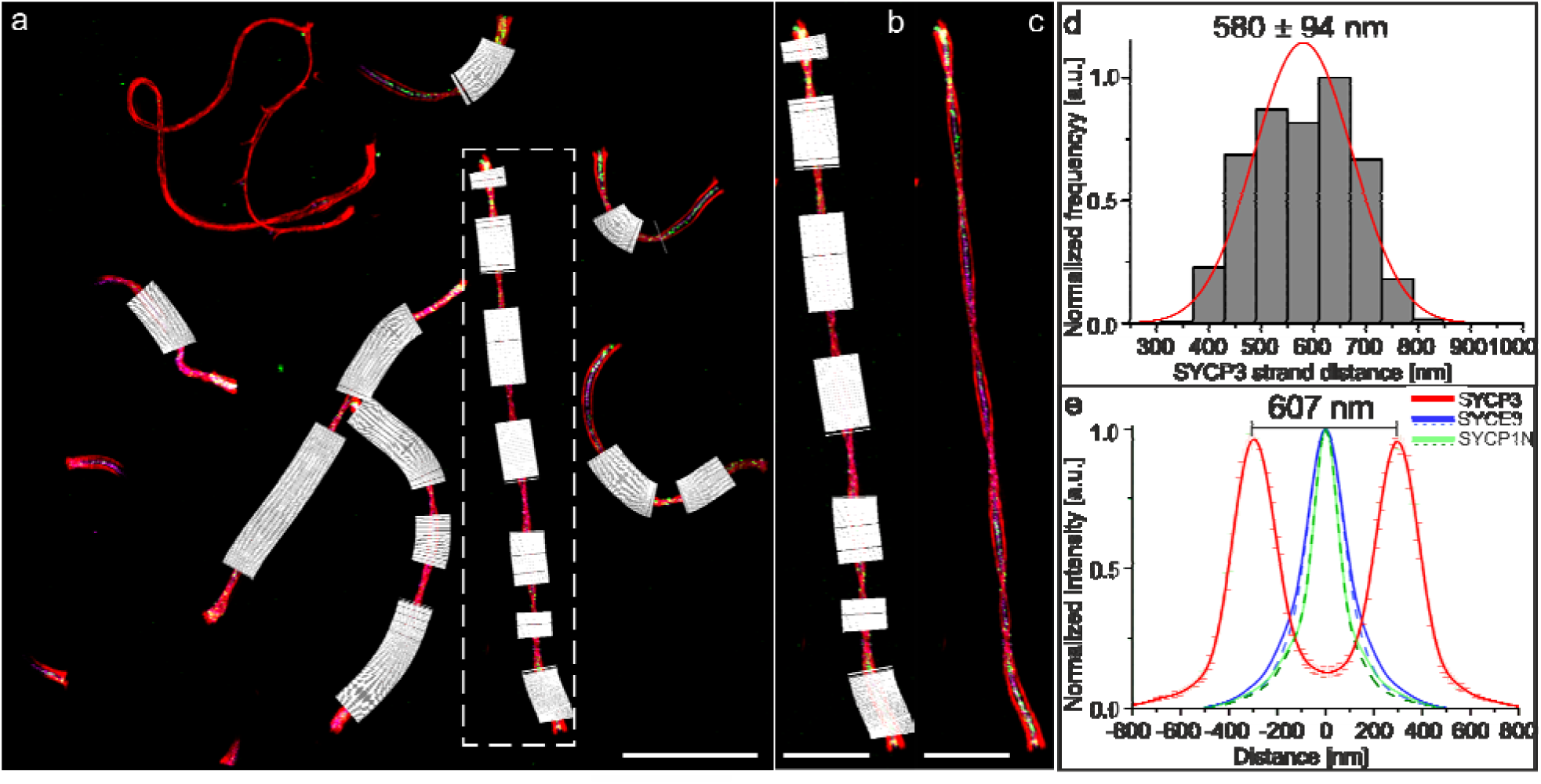
2-channel-mode. **a**, Three-color-SIM image (maximum intensity projection) with line profiles (white) oriented along the SC at regions where SYCP3 shows a bimodal signal distribution using SYCE3 as criterion for the center of the SC. SYCP3 labeled with SeTau647 (red), SYCP1N labeled with Alexa Fluor 488 (green) and SYCE3 labeled with Alexa Fluor 568 (magenta). **b**, Magnified view of white dashed box in (**a**). **c**, Same as (b) without line profiles. **d**, Histogram of SYCP3 distances of 17,607 line profiles determined from 97 MAP-SIM expanded and imaged SCs from two separate experiments. The SYCP3 distance has been determined to 580 ± 93 nm (SD). **e**, Averaged intensity profiles of SYCP3 (red), SYCE3 (blue) and SYCP1N (green) of all analyzed MAP-SIM data in (**d**). Dashed curves show the averaged protein distribution of line profiles only set at frontal views of SYCP3. Whereas solid lines are from data along the complete SC including areas where SYCP3 shows a helical crossing. Fitting of the SYCP3 distribution with a bimodal Gaussian function resulted in a peak-to-peak distance of 607 nm. SYCP1N and SYCE3 showed a monomodal protein distribution of 161.5 ± 1.3 nm (FWHM) for SYCP1N and 229.3 ± 1.2 nm (FWHM) for SYCE3 including SYCP3 crossing point areas. Analysis of the protein distributions without SYCP3 crossing point areas resulted in FWHM of 160.2 ± 1.1 nm for SYCP1N and 207.5 ± 0.78 nm for SYCE3. Scale bar. (**a**) 10 µm. (**b-c**) 5µm.

**Supplementary Figure 2.**
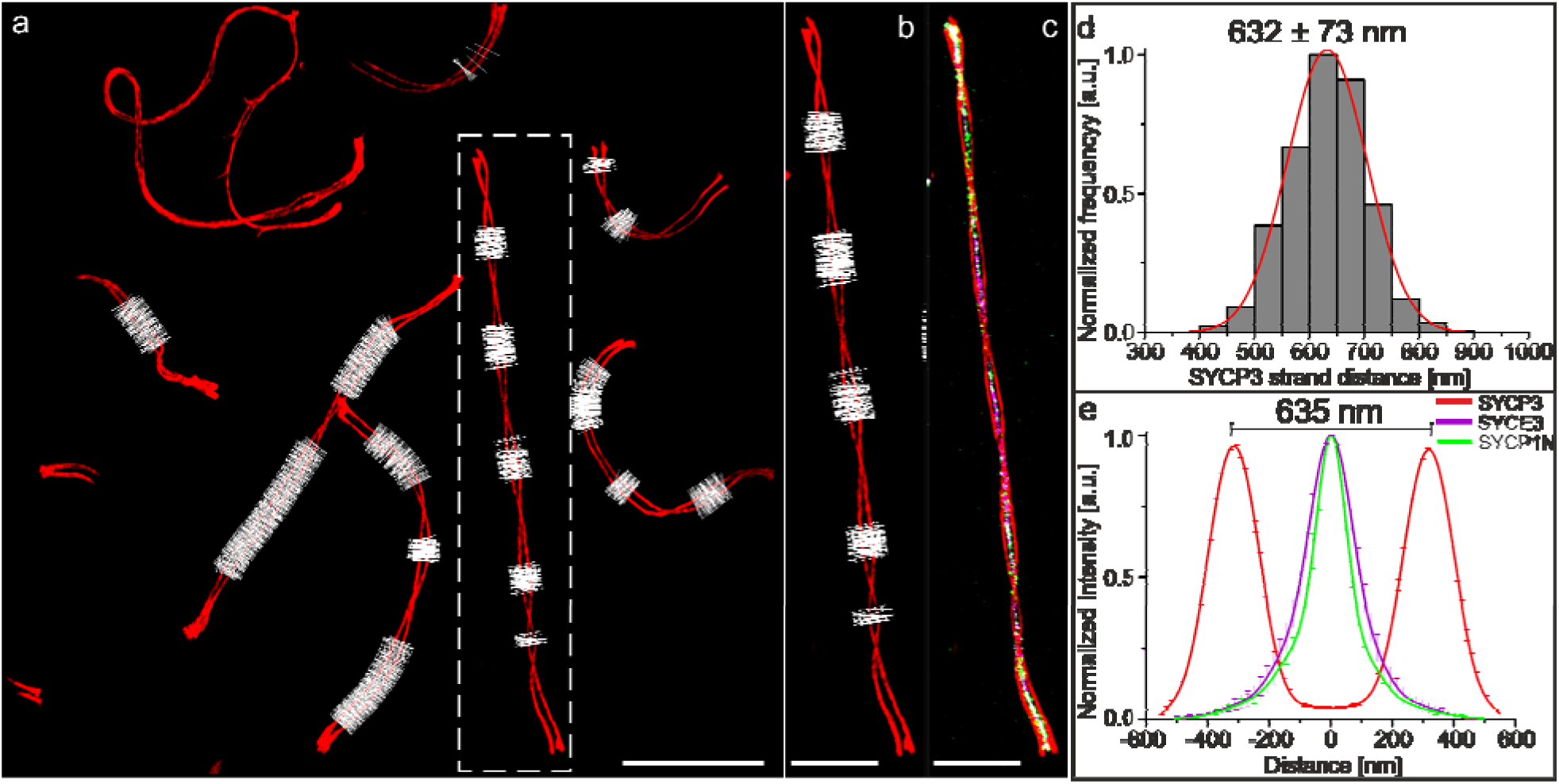
1-channel mode. **a**, Same three-color-SIM image (maximum intensity projection) as shown in Supplementary Fig. 1 but with line profiles (white) oriented along the SC with SYCP3 (red) as criterion for the center of the SC. Profiles are set at regions where a bimodal signal distribution of the protein is occurring. **b**, Magnified view of white dashed box in (**a**). **c**, Same as (b) with SYCP1N (green) and SYCE3 (magenta) without cross-sectional profiles. **d**, Histogram of SYCP3 distances of 18,136 line profiles determined from the same data set analyzed in Supplementary Fig. 1. The SYCP3 distance has been determined to 632 ± 73 nm (SD). Note that different values of the strand distances compared to Supplementary Fig. 1c occur from areas that are not detected for line profiling that are closer to helical crossing overs when using SYCP3 to determine the center of the SC. **e**, Averaged intensity profiles of SYCP3 (red), SYCP1 N (green) and SYCE3 (magenta) of the analyzed MAP-SIM data in (**d**). Fitting of the SYCP3 distribution with a bimodal Gaussian function results in a peak-to-peak distance of 635 nm. Scale bar. (**a**) 10 µm. (**b-c**) 5 µm.

**Supplementary Figure 3.**
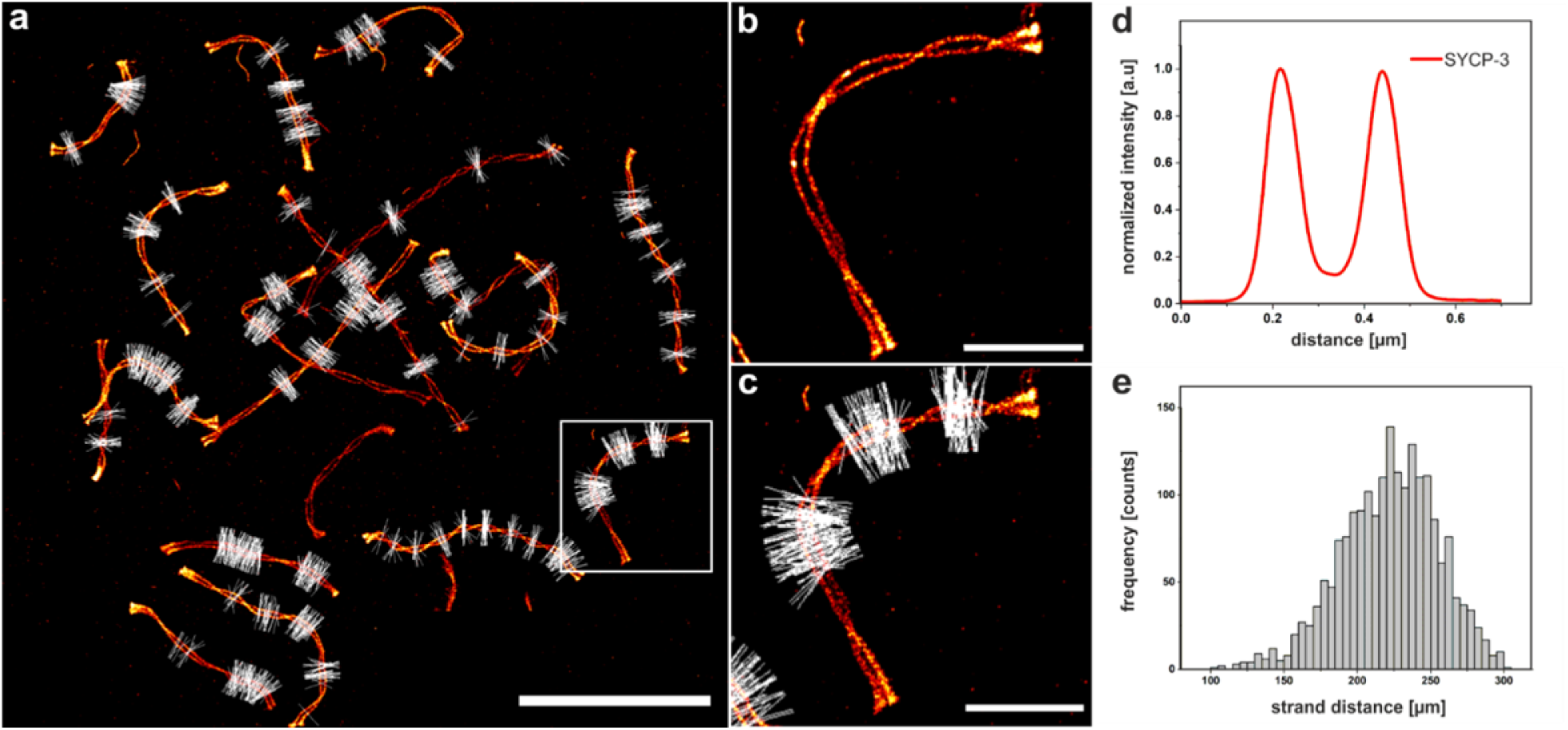
**a**, *d*STORM image of unexpanded SYCP3 labeled with Alexa Fluor 647 (red) with intensity line profiles (white) along the SC using SYCP3 to define the center and orientation of the complexes. **b,** Enlarged view of white box highlighted in (a) without line profiles. **c,** Same as (b) with line profiles (white) along the SYCP3 signal. **d,** Averaged intensity line profile of all profiles shown in (a). A bimodal Gaussian fit of the protein positions gives a peak-to-peak distance of 230 nm (SD). **e,** Histogram of SYCP3 strand distances resulting from line profiles shown in (a) with a mean value of 222 ± 33 nm (SD). Scale bars, (**a**) 10µm, (**b-c**) 2µm.

**Supplementary Figure 4.**
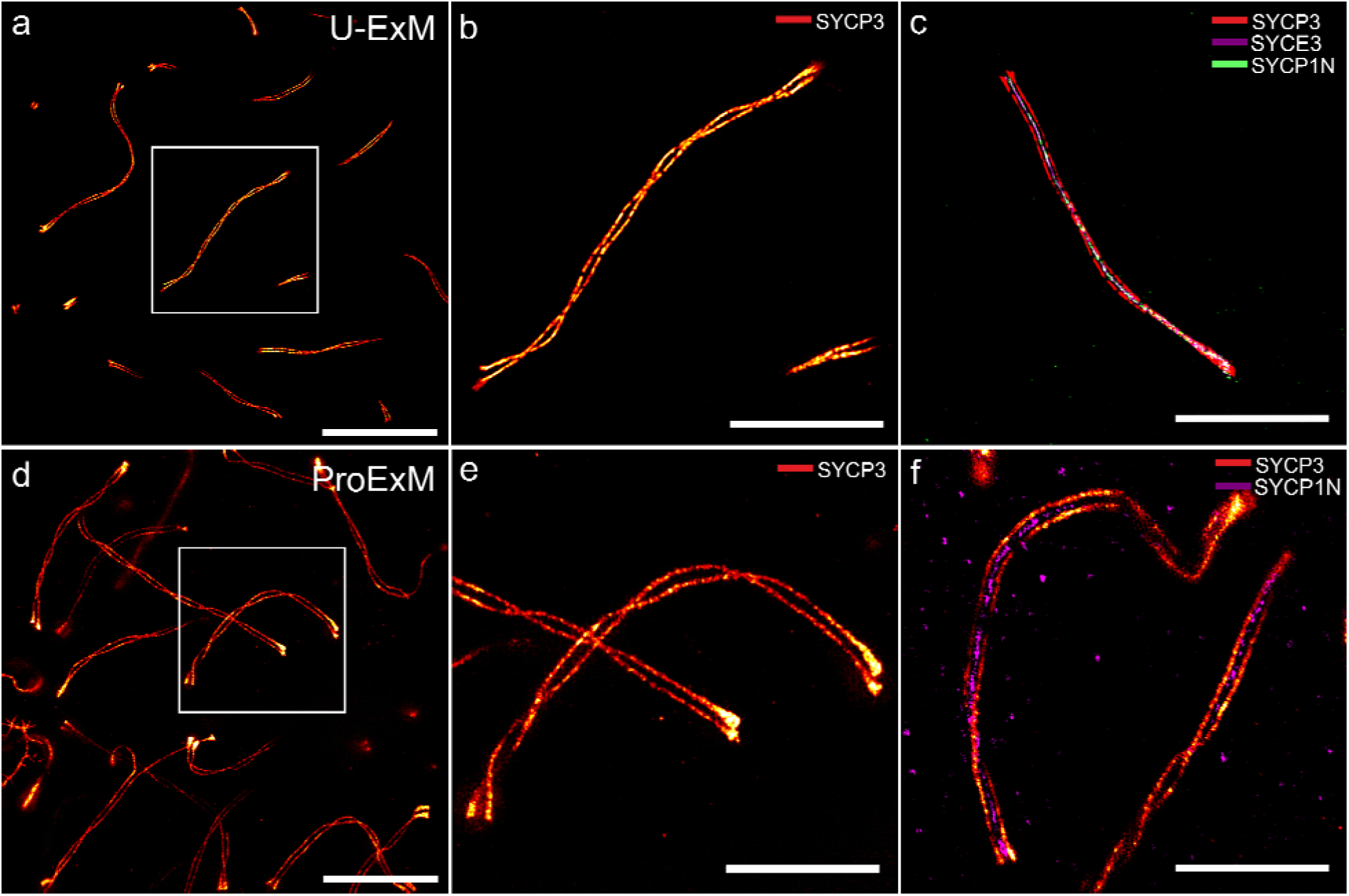
**a**, SIM image (maximum intensity projection of a z-stack) of U-ExM expanded SCs, post-expansion labeled for SYCP3 by immunolabeling with SeTau647. **b**, Magnified view of boxed region in (**a**). **c**, Maximum intensity projection of a z-stack of a multicolor SIM image of an U-ExM expanded SC post-expansion labeled with SeTau647 for SYCP3 (red), SYCE3 labeled with labeled with Alexa Fluor 568 (magenta), and SYCP1N labeled with Alexa Fluor 488 (green). **d**, SIM image (maximum intensity projection of a z-stack) of proExM expanded SCs, pre-expansion labeled for SYCP3 by immunolabeling with Alexa Fluor 488. **e**, Magnified view of boxed region in (**d**). **f**, Maximum intensity projection of a z-stack of a two-color SIM image of a proExM expanded SC pre-expansion labeled with Alexa Fluor 488 for SYCP3 (red) and SYCP1N labeled with labeled with Alexa Fluor 647 (magenta). Scale bars. (**a**) 20 µm, (**b**-**c**) 10 µm, (**d**) 20 µm, (**e**-**f**) 10 µm.

**Supplementary Figure 5.**
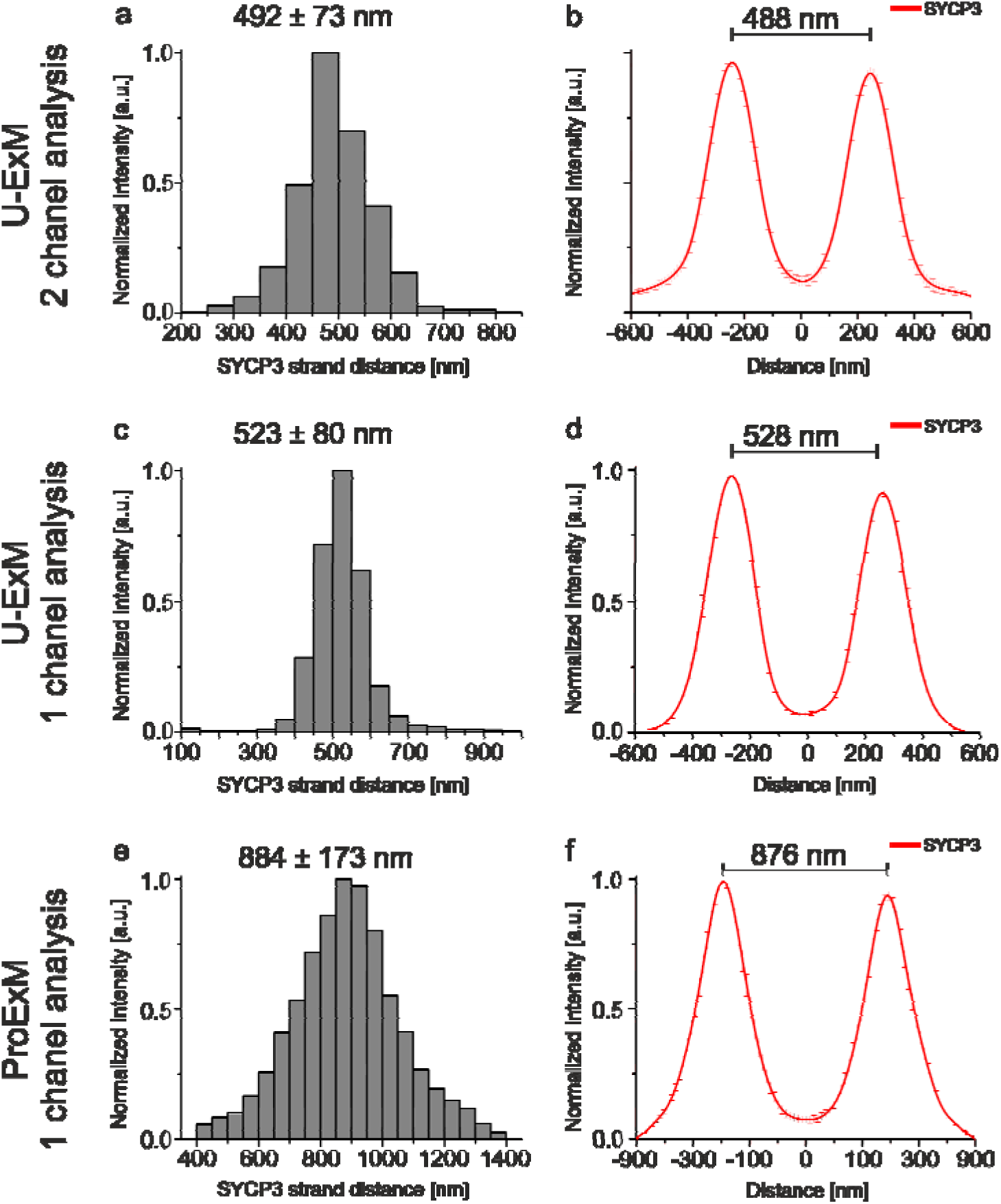
**a**, Histogram of SYCP3 distances determined from U-ExM experiments. The mean distance has been determined to 492 ± 73 nm (SD) from 3,528 intensity line profiles of 24 expanded SCs from two separate expansion experiments. The SYCE3 channel was used to align the line profiles. **b,** Averaged intensity profiles of SYCP3 (red) of U-ExM data from (a). **c-d,** Same data as in (**a**). Here, the SYCP3 channel was used to align the line profiles. The mean distance has been determined to 523 ± 80 nm (SD) from 3528 intensity line profiles of 24 expanded SCs from two separate expansion experiments. **d,** Averaged intensity profiles of SYCP3 (red) of U-ExM data from (c). **e,** Histogram of SYCP3 distances of proExM experiments. The mean distance has been determined to 884 ± 173 nm (SD) from 73427 cross-sectional profiles along 50 expanded SCs. The SYCP3 channel was used to align the line profiles. **f,** Averaged intensity profiles of SYCP3 (red) of proExM data from (e). All data were determined by fitting a bimodal Gaussian function to the histograms. With a bimodal distribution separated by 221nm as measured by *d*STORM (Fig. 2b,e) U-ExM enabled post-expansion labeling with various fluorophores and three-color SIM with an expansion factor of ∼ 2.4x. On the other hand, proExM provided an expansion factor of ∼ 4.0x but pre-expansion labeling resulted generally in a lower labeling density.

**Supplementary Figure 6.**
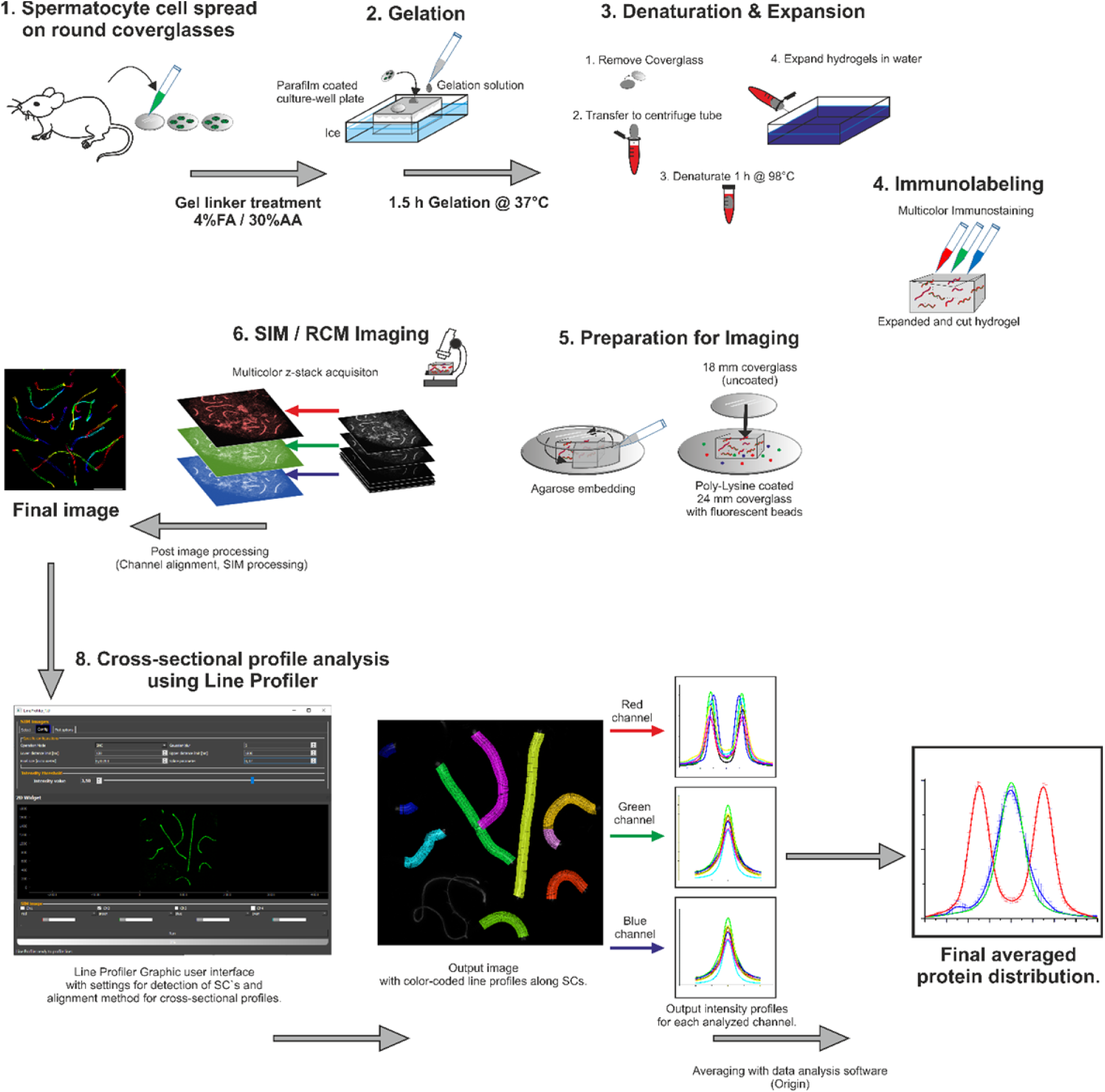
Workflow of MAP-SIM expansion on SCs. **1,** First, spermatocytes are extracted from mice and then spread on round 18 mm coverslips. **2,** After gel linker treatment, cells are gelated on a parafilm coated culture-well on ice. **3,** Hydrogels are then removed carefully from the coverglass and placed into pre-heated denaturation buffer. After 1 hour of denaturation gels are placed into water for expansion. **4,** Samples are immunolabeled successively with primary and secondary antibodies. **5,** Immunolabeled gels are then immobilized on Poly-Lysine coated coverslips and additionally embedded with agarose to prevent drift during imaging. **6**, Samples can then be imaged with SIM or another available imaging technique. After image acquisition post-processing results in the final images (**7**) of SCs. **8,** Images are then further analyzed using the Line Profiler software enabling analysis of protein distributions in several channels. After averaging of cross-sectional profiles a final averaged protein distribution curve is generated.

**Supplementary Video 1. 3D-Multicolor MAP-SIM of SYCP3, SYCP1 N-terminus, and SYCE3.** Expansion microscopy (MAP) of synaptonemal complex (SC) proteins imaged with structured illumination microscopy (SIM). SYCP3 of the lateral element labeled with Setau647 (red), the N-terminus of transverse filament protein SYCP1 labeled with Alexa 488 (green) and SYCE3 of the central element labeled with Alexa 568 (magenta) on a nuclear spreading shown in pachynema. The xy pair can be distinguished by the short synapsed pseudoautosomal region indicated by the presence of all three SC proteins and the larger unsynapsed parts of the x and the y pair that are only associated with SYCP3. The movie sequence shows the progression through the acquired z-stack. SYCP3, SYCP1N and SYCE3 of the triple immunolocalization are further shown separately to provide better visibility of details of the SC’s molecular architecture revealed by MAP-SIM. Note, e.g., that zoomed-in views of the areas where the SC twists (lateral view), suggest a complex architecture of the central element where SYCE3 and the SYCP1 N-terminus reside.

**Supplementary Video 2. Structural details of the SC lateral element revealed by MAP-SIM of SYCP3.** Movie sequence showing the expanded lateral element protein SYCP3 (red, labeled with Setau647) of two SCs in pachynema. Note the fraying of the SYCP3 signal at both ends and the occasional bifurcation of the signal along the length of the SC that is in agreement with EM findings of sub-lateral elements (subLEs) in murine spreadings^38^.

